## Supplemental Tables for "The soil-borne white root rot pathogen *Rosellinia necatrix* expresses antimicrobial proteins during host colonization"

**Supplemental Table 1. Annotation of predicted effector proteins of *R. necatrix* strain R18.**

| **Query ID** | **Accession ID^a^** | **Annotation^b^** | **Organism^c^** | **Query coverage** | **E value** | **Identity (%)^d^** |
| --- | --- | --- | --- | --- | --- | --- |
| **Cell wall degrading enzymes** | | |  |  |  |  |
| FUN_005758 | XM_047973256.1 | 1,4-β-xylosidase | *Xylaria bambusicola* | 100% | 2E-140 | 77.49% |
| FUN_004100 | XM_047973055.1 | 1,4-β-xylosidase | *Xylaria bambusicola* | 99% | 1E-123 | 85.37% |
| FUN_007661 | XM_047977209.1 | 1,4-β-xylosidase | *Xylaria bambusicola* | 99% | 4E-129 | 85.85% |
| FUN_007704 | XM_047978660.1 | Acetylxylan esterase A | *Xylaria bambusicola* | 90% | 0 | 91.73% |
| FUN_003106 | XM_047976713.1 | Carbohydrate esterase family 4 | *Xylaria bambusicola* | 100% | 0 | 78.52% |
| FUN_001755 | XM_047979347.1 | Carbohydrate esterase family 4 | *Xylaria bambusicola* | 100% | 4E-162 | 89.27% |
| FUN_010853 | XM_047976371.1 | Cellulose monooxygenase | *Xylaria bambusicola* | 100% | 4E-131 | 87.32% |
| FUN_008445 | XM_047977156.1 | Cellulose monooxygenase | *Xylaria bambusicola* | 100% | 5E-153 | 87.39% |
| FUN_001148 | XM_046148909.1 | Cutinase | *Microdochium trichocladiopsis* | 100% | 8E-75 | 54.39% |
| FUN_001573 | XM_047972062.1 | Cutinase-like protein | *Xylaria bambusicola* | 91% | 2E-97 | 81.18% |
| FUN_005500 | XM_047972798.1 | Glycoside hydrolase family 10 | *Xylaria bambusicola* | 98% | 0 | 82.68% |
| FUN_000721 | XM_047979386.1 | Glycoside hydrolase family 12 | *Xylaria bambusicola* | 98% | 2E-120 | 73.27% |
| FUN_002439 | XM_047969089.1 | Glycoside hydrolase family 16 | *Xylaria bambusicola* | 100% | 1E-150 | 88.26% |
| FUN_002410 | XM_048004698.1 | Glycoside hydrolase family 16 | *Daldinia vernicosa* | 100% | 2E-110 | 58.33% |
| FUN_010054 | XM_047977697.1 | Glycoside hydrolase family 17 | *Xylaria bambusicola* | 98% | 0 | 85.76% |
| FUN_008369 | XM_047932653.1 | Glycoside hydrolase family 43 | *Daldinia caldariorum* | 100% | 0 | 79.27% |
| FUN_000284 | XM_047979212.1 | Glycoside hydrolase family 43 | *Xylaria bambusicola* | 100% | 0 | 79.88% |
| FUN_008666 | XM_047975058.1 | Glycoside hydrolase family 43 | *Xylaria bambusicola* | 100% | 0 | 82.72% |
| FUN_010499 | XM_047978711.1 | Glycoside hydrolase family 45 | *Xylaria bambusicola* | 99% | 2E-133 | 85.24% |
| FUN_007581 | XM_047971870.1 | Glycoside hydrolase family 5 | *Xylaria bambusicola* | 99% | 0 | 84.94% |
| FUN_004897 | XM_047976082.1 | Lytic polysaccharide monooxygenase | *Xylaria bambusicola* | 100% | 3E-126 | 84.11% |
| FUN_007011 | XM_022614464.1 | Mutanase | *Colletotrichum orchidophilum* | 98% | 3E-48 | 70.00% |
| FUN_008395 | XM_047972336.1 | O-acetylxylan esterase | *Xylaria bambusicola* | 100% | 3E-141 | 89.33% |
| FUN_007837 | XM_040856674.1 | Pectate lyase | *Pseudomassariella vexata* | 100% | 6E-129 | 84.11% |
| FUN_004715 | XM_047972211.1 | Polysaccharide lyase | *Xylaria bambusicola* | 100% | 4E-154 | 87.71% |
| FUN_001279 | XM_047975706.1 | Polysaccharide lyase family 7 protein | *Xylaria bambusicola* | 100% | 4E-133 | 82.38% |
| FUN_009124 | XM_047969579.1 | Putative endo-1,3(4)-beta-glucanase | *Xylaria bambusicola* | 99% | 2E-142 | 78.03% |
| FUN_010529 | XM_047974852.1 | Xylanase | *Xylaria bambusicola* | 100% | 7E-129 | 91.41% |
| FUN_009151 | XM_047976406.1 | Lysozyme-like domain-containing protein | *Xylaria bambusicola* | 99% | 1E-95 | 83.73% |
| FUN_003275 | XM_018318865.1 | Muramidase | *Purpureocillium lilacinum* | 92% | 2E-97 | 64.29% |
| FUN_005762 | XM_047972660.1 | Putative muramidase | *Xylaria bambusicola* | 99% | 1E-102 | 71.16% |
| FUN_001575 | XM_047978226.1 | Chloroperoxidase | *Xylaria bambusicola* | 99% | 2E-133 | 76.17% |
| **Carbohydrate binding** | |  |  |  |  |  |
| FUN_003047 | XM_047979309.1 | Concanavalin A-like lectin | *Xylaria bambusicola* | 99% | 1E-176 | 81.06% |
| FUN_003763 | XM_047969123.1 | WSC domain-containing protein | *Xylaria bambusicola* | 88% | 3E-121 | 77.39% |
| FUN_000265 | XM_047976926.1 | WSC domain-containing protein | *Xylaria bambusicola* | 100% | 0 | 79.34% |
| **Lysin motif (LysM) effectors** | |  |  |  |  |  |
| FUN_000206 | XM_001268710.1 | LysM domain-containing protein | *Aspergillus clavatus* | 83% | 4E-08 | 38.46% |
| FUN_006904 | XM_001268710.1 | LysM domain-containing protein | *Aspesgillus clavatus* | 98% | 8E-25 | 37.84% |
| **Hydrophobins** | |  | | | | |
| FUN_007007 | XM_040863415.1 | Cerato-ulmin family protein | *Pseudomassariella vexata* | 81% | 1E-32 | 76.71% |
| FUN_000192 | XM_047977015.1 | Cerato-ulmin family protein | *Xylaria bambusicola* | 95% | 5E-40 | 82.28% |
| FUN_001605 | XM_047978428.1 | Hydrophobin | *Xylaria bambusicola* | 100% | 1E-31 | 57.55% |
| FUN_011531 | XM_047976151.1 | Hydrophobin-like protein | *Xylaria bambusicola* | 84% | 6E-40 | 90.14% |
| **Proteases** |  |  |  |  |  |  |
| FUN_001370 | XM_047978948.1 | Acid protease | *Xylaria bambusicola* | 100% | 7E-106 | 65.82% |
| FUN_004271 | XM_036641540.1 | Neutral protease 2-like protein | *Colletotrichum siamense* | 99% | 1E-114 | 51.81% |
| FUN_000625 | XM_033580201.1 | Papain inhibitor | *Daldinia childiae* | 97% | 2E-36 | 65.12% |
| **Toxins** | | | |  |  |  |
| FUN_002843 | XM_047969234.1 | Necrosis inducing protein-domain-containing protein | *Xylaria bambusicola* | 100% | 6E-147 | 82.45% |
| FUN_005483 | XM_046238898.1 | Necrosis inducing protein-domain-containing protein | *Ilyonectria robusta* | 92% | 3E-82 | 54.20% |
| FUN_010366 | XM_047972065.1 | Cerato-platanin family protein | *Xylaria bambusicola* | 100% | 2E-64 | 80.83% |
| **Others** |  |  |  |  |  |  |
| FUN_000283 | XM_047980145.1 | Deoxyribonuclease NucA/NucB | *Xylaria bambusicola* | 99% | 1E-124 | 81.73% |
| FUN_003451 | XM_047976696.1 | Guanine-specific ribonuclease N1 | *Xylaria bambusicola* | 100% | 3E-61 | 79.82% |
| FUN_009264 | XM_047973582.1 | emp24/gp25L/p24 family/GOLD-domain-containing protein | *Xylaria bambusicola* | 100% | 9E-108 | 75.37% |
| FUN_009552 | XM_047973428.1 | emp24/gp25L/p24 family/GOLD-domain-containing protein | *Xylaria bambusicola* | 100% | 4E-127 | 88.61% |
| FUN_005699 | XM_047975004.1 | emp24/gp25L/p24 family/GOLD-domain-containing protein | *Xylaria bambusicola* | 100% | 4E-123 | 90.96% |
| FUN_001432 | XM_047970391.1 | 2OG-Fe(II) oxygenase superfamily protein | *Xylaria bambusicola* | 100% | 5E-133 | 75.95% |
| FUN_011359 | XM_048011353.1 | FMN-dependent alpha-hydroxy acid dehydrogenase | *Daldinia vernicosa* | 95% | 2E-69 | 74.10% |
| FUN_011399 | KY782142.1 | Heme-thiolate peroxidase | *Ustulina deusta* | 99% | 1E-124 | 73.33% |
| FUN_001799 | XM_047970313.1 | Intradiol ring-cleavage dioxygenase | *Xylaria bambusicola* | 100% | 0 | 74.93% |
| FUN_009516 | XM_049306200.1 | Putative choline dehydrogenase | *Daldinia loculata* | 87% | 3E-28 | 63.10% |
| FUN_004320 | XM_047974920.1 | ERV/ALR sulfhydryl oxidase | *Xylaria bambusicola* | 91% | 1E-87 | 79.49% |
| FUN_002834 | XM_047975475.1 | Acyl transferase | *Xylaria bambusicola* | 76% | 1E-93 | 78.57% |
| FUN_006056 | XM_047978164.1 | Apc13p ubiquitin ligase | *Xylaria bambusicola* | 87% | 4E-59 | 65.41% |
| FUN_005907 | XM_047975260.1 | Cyanovirin-N | *Xylaria bambusicola* | 93% | 1E-57 | 66.94% |
| FUN_006163 | XM_046160209.1 | Cyclophilin-like protein | *Microdochium trichocladiopsis* | 100% | 1E-102 | 81.67% |
| **Unknown** |  |  |  |  |  |  |
| FUN_010039 | XM_033580564.1 | Bys1 domain-containing protein | *Daldinia childiae* | 100% | 5E-79 | 82.09% |
| FUN_005325 | XM_047969535.1 | Met-10+ like-protein-domain-containing protein | *Xylaria bambusicola* | 100% | 1E-95 | 70.79% |

**^a^**Database accession ID where subject was deposited.

**^b^**Annotation of the best hit using BLAST (tblastn).

**^c^**Organism where the homolog was annotated.

**^d^**Percentage of identity of the query.

**Supplemental Table 2.** **Single nucleotide polymorphism ratios (SNPs in %) and presence-absence variation for predicted effector genes in the *R. necatrix* strains sequenced in this study using *R. necatrix* strain R18 as a reference.**

| **Effector in *R. necatrix* strain R18 (gene ID)** | **Homologs in other *R. necatrix* strains^*^** | | | | | | | |
| --- | --- | --- | --- | --- | --- | --- | --- | --- |
|  | **Rn19** | **CH12** | **Rn400** | **R10** | **R25** | **R27** | **R28** | **R30** |
| FUN_000053 | 3.29 | 3.29 | 4.15 | 0.00 | 0.17 | 0.00 | 0.00 | 0.17 |
| FUN_000192 | 3.63 | 3.29 | 3.29 | 0.00 | 0.17 | 0.00 | 0.00 | 0.17 |
| FUN_000206 | 3.98 | 3.81 | 3.81 | 0.17 | 0.17 | 0.17 | 0.00 | 0.17 |
| FUN_000251 | 3.98 | 3.98 | 3.98 | 0.00 | 0.35 | 0.87 | 0.87 | 0.35 |
| FUN_000265 | 8.82 | 8.65 | 8.82 | 0.00 | 1.04 | 1.21 | 1.21 | 1.04 |
| FUN_000283 | 6.06 | 6.06 | 5.36 | A | 1.56 | 1.56 | 1.38 | 1.56 |
| FUN_000284 | 7.61 | 7.61 | 8.13 | 0.35 | 1.04 | 1.56 | 1.56 | 1.04 |
| FUN_000359 | 3.46 | 4.50 | 3.11 | 0.00 | 1.90 | 1.56 | 1.56 | 1.56 |
| FUN_000367 | 2.25 | 2.25 | 2.25 | 0.00 | 0.17 | 0.00 | 0.00 | 0.17 |
| FUN_000625 | 2.60 | 2.60 | 2.60 | 0.00 | 0.69 | 0.00 | 0.00 | 0.69 |
| FUN_000721 | 3.81 | 3.81 | 1.21 | 0.00 | 1.04 | 0.00 | 0.17 | 1.04 |
| FUN_000856 | 2.42 | 2.60 | 2.60 | 0.17 | 0.87 | 0.35 | 0.35 | 1.04 |
| FUN_000873 | 2.77 | 2.77 | 2.77 | 0.00 | 0.87 | 0.87 | 0.87 | 0.87 |
| FUN_000886 | 4.33 | 7.09 | 4.33 | 0.00 | 1.38 | 1.21 | 1.21 | 1.56 |
| FUN_000960 | 7.61 | 7.61 | 8.65 | 0.52 | 2.08 | 1.73 | 1.38 | 1.90 |
| FUN_001031 | 2.42 | 3.29 | 2.42 | 0.17 | 0.69 | 2.94 | 3.29 | 0.69 |
| FUN_001148 | 3.29 | 3.29 | 3.29 | 0.00 | 1.21 | 0.52 | 0.52 | 1.04 |
| FUN_001231 | 2.25 | 2.25 | 2.42 | 0.00 | 0.17 | 0.00 | 0.00 | 0.17 |
| FUN_001279 | 6.75 | 6.75 | 6.75 | 0.00 | 1.73 | 1.73 | 0.87 | 1.73 |
| FUN_001370 | 8.13 | 7.61 | 7.96 | 0.00 | 2.08 | 2.08 | 0.87 | 2.08 |
| FUN_001432 | 5.54 | 5.19 | 5.19 | 0.00 | 1.38 | 1.90 | 0.87 | 1.38 |
| FUN_001497 | 2.42 | 2.60 | 2.77 | 0.00 | 0.00 | 0.35 | 0.69 | 0.00 |
| FUN_001504 | 3.29 | 3.11 | 3.29 | 0.00 | 0.00 | 0.00 | 0.00 | 0.00 |
| FUN_001515 | 2.08 | 2.08 | 2.08 | 0.00 | 0.69 | 0.00 | 0.00 | 0.87 |
| FUN_001527 | 5.88 | 6.23 | 5.88 | 3.98 | 1.38 | 6.06 | 0.00 | 6.23 |
| FUN_001573 | 4.50 | 4.50 | 4.50 | 0.00 | A | 0.00 | 0.00 | 0.69 |
| FUN_001575 | 6.06 | 5.88 | 5.88 | 0.00 | 1.38 | 0.00 | 0.00 | 1.38 |
| FUN_001585 | 4.67 | 4.84 | 3.98 | 0.00 | 0.87 | 0.17 | 0.00 | 0.69 |
| FUN_001595 | 2.08 | 2.25 | 2.25 | 0.52 | 0.52 | 0.00 | 0.00 | 0.52 |
| FUN_001605 | 9.86 | 9.86 | 9.86 | 0.00 | 3.11 | 0.00 | 0.00 | 2.94 |
| FUN_001636 | 2.94 | 2.94 | 2.77 | 0.00 | 0.00 | 0.00 | 0.00 | 0.00 |
| FUN_001689 | 6.57 | 6.57 | 6.92 | 0.17 | 2.60 | 0.00 | 0.00 | 2.60 |
| FUN_001755 | 9.17 | 3.46 | 3.29 | 0.00 | 1.04 | 0.69 | 0.00 | 0.87 |
| FUN_001798 | 9.69 | 9.52 | 9.17 | 0.00 | 0.00 | 0.00 | 0.00 | 0.00 |
| FUN_001799 | 1.56 | 0.52 | 0.87 | 0.17 | 0.17 | 0.00 | 0.00 | 0.00 |
| FUN_001917 | 2.42 | 2.42 | 1.90 | 0.00 | 0.52 | 0.00 | 0.00 | 0.52 |
| FUN_002005 | 4.84 | 4.67 | 4.15 | 0.17 | 0.35 | 2.08 | 2.08 | 0.35 |
| FUN_002053 | 1.04 | 1.04 | 1.04 | 0.00 | 0.17 | 0.35 | 0.17 | 0.35 |
| FUN_002114 | 3.46 | 3.29 | 3.29 | 0.00 | 0.87 | 0.35 | 0.35 | 0.87 |
| FUN_002120 | 4.50 | 4.50 | 3.46 | 0.00 | 0.87 | 1.21 | 1.21 | 0.87 |
| FUN_002121 | 4.50 | 4.50 | 3.46 | 0.00 | 0.00 | 0.00 | 1.21 | 0.00 |
| FUN_002234 | 3.11 | 2.60 | 2.77 | 0.00 | 0.35 | 0.52 | 0.52 | 0.35 |
| FUN_002337 | 8.13 | 8.13 | 8.13 | 8.30 | 0.52 | 0.00 | 0.00 | 0.52 |
| FUN_002389 | 0.69 | 0.69 | 0.87 | 0.35 | 0.52 | 0.69 | 0.52 | 0.52 |
| FUN_002410 | 5.88 | 5.88 | 5.88 | 0.00 | 2.08 | 1.90 | 1.90 | 2.08 |
| FUN_002439 | 11.76 | 11.76 | 11.76 | 0.00 | 2.77 | 2.60 | 0.00 | 2.77 |
| FUN_002458 | 2.42 | 2.42 | 2.42 | 0.17 | 0.35 | 1.56 | 0.35 | 0.35 |
| FUN_002489 | 4.33 | 5.02 | 5.02 | 0.17 | 1.56 | 0.87 | 1.04 | 1.56 |
| FUN_002758 | 1.21 | 1.38 | 1.38 | 0.00 | 0.17 | 1.04 | 1.04 | 0.17 |
| FUN_002834 | 3.11 | 2.94 | 3.29 | 0.00 | 0.69 | 0.69 | 0.69 | 0.69 |
| FUN_002843 | 5.71 | 5.71 | 5.71 | 0.00 | 0.52 | 1.04 | 1.04 | 0.52 |
| FUN_003003 | 1.38 | 1.38 | 1.38 | 0.00 | 0.17 | 0.00 | 0.00 | 0.17 |
| FUN_003018 | 1.38 | 0.69 | 0.69 | 0.00 | 0.00 | 0.00 | 0.00 | 0.00 |
| FUN_003047 | 4.50 | A | 4.33 | 0.00 | 0.00 | 0.00 | 0.00 | 0.00 |
| FUN_003077 | 2.77 | 2.77 | 3.29 | 0.00 | 0.17 | 0.35 | 0.35 | 0.17 |
| FUN_003106 | 9.52 | 9.69 | 9.52 | 0.00 | 0.00 | 0.87 | 0.87 | 0.00 |
| FUN_003133 | 2.77 | 2.25 | 1.90 | 0.52 | 1.90 | 0.00 | 0.52 | 1.90 |
| FUN_003175 | 3.81 | 3.63 | 3.81 | 0.35 | 0.17 | 0.17 | 0.17 | 0.17 |
| FUN_003209 | 3.63 | 3.46 | 3.29 | 0.00 | 0.00 | 0.00 | 0.00 | 1.21 |
| FUN_003212 | 4.67 | 4.67 | 4.67 | 0.00 | 0.35 | 0.35 | 0.35 | 3.46 |
| FUN_003275 | 2.94 | 2.77 | 2.77 | 0.00 | 0.00 | 0.00 | 0.00 | 0.00 |
| FUN_003286 | 3.46 | 3.46 | 3.46 | 0.00 | 0.00 | 0.00 | 0.00 | 0.00 |
| FUN_003451 | 3.63 | 3.63 | 3.63 | 0.00 | 0.35 | 0.52 | 0.52 | 0.35 |
| FUN_003513 | 1.38 | 1.38 | 1.38 | 0.00 | 0.00 | 0.35 | 0.35 | 0.00 |
| FUN_003763 | 4.67 | 4.67 | 4.67 | 0.35 | 0.17 | 0.17 | 0.00 | 0.17 |
| FUN_003889 | 8.13 | 8.30 | 8.13 | 0.00 | 2.94 | 0.00 | 0.00 | 2.94 |
| FUN_003947 | 9.17 | 8.82 | 8.82 | 0.00 | 0.69 | 0.00 | 0.00 | 0.87 |
| FUN_004082 | 4.15 | 4.15 | 4.15 | 0.17 | 0.00 | 0.00 | 0.00 | 0.00 |
| FUN_004100 | 5.54 | 5.54 | 5.88 | 0.00 | 0.35 | 0.52 | 0.52 | 0.35 |
| FUN_004157 | 4.67 | 1.21 | 6.40 | 0.52 | 7.96 | 7.96 | 0.00 | 4.84 |
| FUN_004244 | 5.36 | 7.44 | 4.50 | 0.00 | 1.38 | 0.00 | 0.00 | 1.38 |
| FUN_004254 | 1.73 | 1.90 | 1.73 | 0.35 | 0.00 | 0.35 | 0.00 | 0.17 |
| FUN_004271 | 3.63 | 3.63 | 3.46 | 0.00 | 0.17 | 0.00 | 0.00 | 0.17 |
| FUN_004305 | 5.71 | 5.88 | 5.54 | 0.17 | 0.00 | 0.87 | 0.69 | 0.00 |
| FUN_004320 | 3.46 | 3.46 | 3.46 | 0.00 | 1.04 | 1.38 | 1.38 | 1.04 |
| FUN_004483 | 4.15 | 4.15 | 4.15 | 0.00 | 0.00 | 1.04 | 1.04 | 0.00 |
| FUN_004569 | 2.42 | 2.77 | 2.77 | 0.00 | 2.60 | 2.08 | 1.90 | 2.60 |
| FUN_004715 | 1.90 | 1.73 | 1.90 | 0.00 | 0.35 | 0.35 | 0.35 | 0.35 |
| FUN_004770 | 4.15 | 4.15 | 4.15 | 0.00 | 0.00 | 0.00 | 0.00 | 0.00 |
| FUN_004897 | 4.67 | 4.33 | 4.50 | 0.17 | 0.00 | 0.00 | 0.00 | 0.00 |
| FUN_004961 | 4.15 | 4.15 | 4.33 | 0.00 | 0.35 | 0.17 | 0.00 | 0.35 |
| FUN_005189 | 5.36 | 1.38 | 5.19 | 0.00 | 0.00 | 0.69 | 0.87 | 0.00 |
| FUN_005222 | 4.84 | 4.67 | 4.67 | 0.00 | 0.00 | 0.00 | 0.00 | 0.00 |
| FUN_005300 | 4.67 | 1.90 | 5.36 | 0.00 | 1.21 | 1.21 | 1.21 | 1.21 |
| FUN_005325 | 0.87 | 1.38 | 1.38 | 0.00 | 0.35 | 1.04 | 1.04 | 0.35 |
| FUN_005445 | 1.56 | 1.90 | 1.73 | 0.00 | 0.17 | 0.00 | 0.00 | 0.17 |
| FUN_005483 | 2.77 | 4.84 | 5.02 | 0.69 | 2.94 | 2.94 | 2.08 | 2.60 |
| FUN_005500 | 4.67 | 3.46 | 6.06 | 0.69 | 3.46 | 1.90 | 2.08 | 3.81 |
| FUN_005538 | 4.33 | 3.63 | 3.63 | 0.17 | 1.04 | 1.21 | 1.21 | 1.04 |
| FUN_005543 | 5.54 | 0.00 | 5.02 | 0.00 | 1.21 | 0.00 | 0.00 | 0.00 |
| FUN_005544 | 7.27 | 7.27 | 7.96 | 0.00 | 0.35 | 0.87 | 0.87 | 0.35 |
| FUN_005699 | 3.46 | 2.94 | 2.94 | 0.00 | 0.17 | 0.00 | 0.00 | 0.17 |
| FUN_005740 | 1.21 | 1.21 | 1.21 | 0.00 | 0.35 | 0.00 | 0.00 | 0.35 |
| FUN_005758 | 6.75 | 4.33 | 4.33 | 0.17 | 1.90 | 4.15 | 0.17 | 4.15 |
| FUN_005762 | 3.29 | 3.46 | 3.29 | 0.00 | 0.17 | 0.00 | 0.17 | 0.17 |
| FUN_005785 | 4.50 | 4.50 | 4.33 | 0.00 | 0.35 | 0.00 | 0.00 | 0.35 |
| FUN_005787 | 3.63 | 3.63 | 3.63 | 0.00 | 0.35 | 0.00 | 0.00 | 0.35 |
| FUN_005834 | 3.11 | 3.11 | 3.11 | 0.00 | 0.69 | 0.00 | 0.00 | 0.69 |
| FUN_005847 | 8.30 | 8.30 | 8.13 | 0.00 | 0.00 | 0.00 | 0.00 | 0.00 |
| FUN_005850 | 4.15 | 4.84 | 3.46 | 0.00 | 0.52 | 0.00 | 0.00 | 0.52 |
| FUN_005883 | 4.33 | 4.50 | 4.67 | 0.00 | 0.87 | 0.87 | 0.87 | 0.87 |
| FUN_005907 | 4.33 | 4.15 | 4.33 | 0.00 | 0.52 | 0.00 | 0.00 | 0.52 |
| FUN_005940 | 5.19 | 5.19 | 4.67 | 0.00 | 0.00 | 2.60 | 2.60 | 0.00 |
| FUN_005952 | 4.84 | 3.98 | 4.33 | 0.00 | 0.00 | 0.69 | 0.69 | 0.00 |
| FUN_005989 | 5.88 | 6.57 | 6.06 | 0.00 | 1.04 | 0.17 | 0.17 | 0.87 |
| FUN_005998 | 6.06 | 6.23 | 5.88 | 0.00 | 0.35 | 0.17 | 0.00 | 0.35 |
| FUN_006007 | 8.13 | 8.30 | 7.61 | 0.00 | 0.00 | 2.77 | 2.77 | 0.00 |
| FUN_006010 | 4.15 | 4.33 | 3.81 | 0.00 | 0.17 | 0.35 | 0.35 | 0.17 |
| FUN_006056 | 3.63 | 3.63 | 3.81 | 0.00 | 0.00 | 0.17 | 0.00 | 0.00 |
| FUN_006163 | 3.98 | 1.90 | 4.15 | 0.00 | 0.17 | 1.90 | 1.90 | 0.17 |
| FUN_006164 | 3.98 | 1.90 | A | 0.00 | 0.00 | 1.73 | 0.00 | 0.17 |
| FUN_006190 | 5.36 | 5.88 | 5.71 | 0.00 | 6.06 | 0.69 | 6.23 | 0.00 |
| FUN_006412 | 6.40 | 4.67 | 4.67 | 0.00 | 1.04 | 0.52 | 1.04 | 1.04 |
| FUN_006481 | 3.29 | 2.08 | 3.29 | 0.00 | 0.17 | 0.52 | 0.52 | 0.17 |
| FUN_006483 | 3.98 | 4.33 | 4.67 | 0.17 | 0.17 | 0.52 | 0.35 | 0.17 |
| FUN_006775 | 8.65 | 8.65 | 8.65 | 0.00 | 0.87 | 0.35 | 0.35 | 0.87 |
| FUN_006835 | 2.42 | 2.42 | 2.77 | 0.00 | 1.04 | 1.56 | 1.38 | 1.04 |
| FUN_006841 | 5.36 | 4.84 | 5.36 | 0.00 | 0.52 | 0.87 | 0.87 | 0.52 |
| FUN_006904 | 6.23 | 6.23 | 7.09 | 0.00 | 0.69 | 0.00 | 0.00 | 0.69 |
| FUN_006905 | 4.67 | 4.50 | 6.06 | 0.00 | 0.00 | 0.17 | 0.00 | 0.00 |
| FUN_007007 | 5.88 | 5.88 | 7.27 | 0.00 | 1.38 | 0.00 | 0.00 | 1.38 |
| FUN_007011 | 2.42 | 2.60 | 1.73 | 0.00 | 0.17 | 0.00 | 0.00 | 0.17 |
| FUN_007012 | 6.06 | 5.71 | 6.23 | 0.00 | 0.35 | 0.00 | 0.00 | 0.35 |
| FUN_007129 | 4.84 | 5.36 | 5.36 | 0.00 | 0.52 | 0.00 | 0.00 | 0.52 |
| FUN_007254 | 4.33 | 4.33 | 4.50 | 0.00 | 0.87 | 1.04 | 1.04 | 0.87 |
| FUN_007288 | 8.13 | 7.27 | 8.13 | 0.00 | 1.38 | 0.87 | 0.87 | 1.38 |
| FUN_007567 | 2.77 | 2.94 | 2.60 | 0.00 | 0.00 | 0.17 | 0.17 | 0.00 |
| FUN_007581 | 7.44 | 7.27 | 6.92 | 0.00 | 0.35 | 0.52 | 0.52 | 0.35 |
| FUN_007661 | 4.67 | 3.98 | 4.15 | 0.00 | 0.69 | 0.87 | 0.87 | 0.69 |
| FUN_007684 | 6.75 | 7.27 | 6.57 | 0.00 | 0.00 | 1.38 | 1.38 | 0.17 |
| FUN_007704 | 6.23 | 6.40 | 5.71 | 0.00 | 0.00 | 0.52 | 0.69 | 0.00 |
| FUN_007825 | 2.42 | 2.42 | 2.42 | 0.00 | 0.00 | 1.56 | 1.56 | 0.00 |
| FUN_007837 | 4.84 | 1.73 | 5.02 | 0.17 | 0.69 | 1.04 | 1.21 | 0.69 |
| FUN_007838 | 1.21 | 1.04 | 1.21 | 0.00 | 0.00 | 0.35 | 0.17 | 0.00 |
| FUN_007850 | 2.08 | 1.21 | 2.08 | 0.00 | 0.00 | 0.00 | 0.00 | 0.35 |
| FUN_007851 | 3.63 | 2.08 | 3.63 | 0.00 | 0.52 | 0.17 | 0.17 | 0.52 |
| FUN_007855 | 5.19 | 3.63 | 5.71 | 0.00 | 0.69 | 0.52 | 0.52 | 0.69 |
| FUN_007860 | 6.40 | 5.19 | 6.40 | 0.00 | 2.25 | 1.73 | 1.56 | 2.25 |
| FUN_007991 | 1.56 | 6.40 | 1.73 | 0.17 | 0.69 | 0.35 | 0.35 | 0.69 |
| FUN_008329 | 2.60 | 2.60 | 2.60 | 0.00 | 1.38 | 0.00 | 0.00 | 1.38 |
| FUN_008352 | 2.42 | 2.42 | 2.42 | 0.00 | 1.04 | 0.00 | 0.00 | 1.04 |
| FUN_008369 | 6.40 | 6.23 | 6.40 | 0.00 | 0.69 | 0.00 | 0.00 | 0.69 |
| FUN_008395 | 3.63 | 2.94 | 3.63 | 0.00 | 1.21 | 0.00 | 0.00 | 1.21 |
| FUN_008445 | 2.25 | 2.60 | 2.60 | 0.00 | 0.35 | 0.00 | 0.00 | 0.35 |
| FUN_008500 | 2.60 | 1.90 | 2.08 | 0.00 | 0.17 | 0.00 | 0.00 | 0.17 |
| FUN_008562 | 2.77 | 3.11 | 2.25 | 0.17 | 0.00 | 0.17 | 0.00 | 0.00 |
| FUN_008574 | 5.02 | 5.71 | 4.84 | 0.17 | 2.60 | 0.00 | 0.00 | 2.42 |
| FUN_008666 | 5.02 | 5.71 | 5.54 | 0.00 | 0.35 | 0.00 | 0.17 | 0.35 |
| FUN_008681 | A | A | A | 0.00 | 0.52 | 0.52 | 0.69 | 0.52 |
| FUN_008714 | 6.57 | 3.98 | 3.98 | 0.00 | 1.04 | 1.04 | 1.04 | 1.04 |
| FUN_008830 | 5.02 | 5.19 | 5.19 | 0.00 | 0.52 | 0.52 | 0.52 | 0.52 |
| FUN_009063 | 6.57 | 6.57 | 6.57 | 0.00 | 2.42 | 0.00 | 0.17 | 2.42 |
| FUN_009073 | 5.88 | 5.19 | 5.19 | 0.17 | 0.87 | 0.00 | 0.17 | 0.87 |
| FUN_009124 | 4.33 | 4.33 | 4.33 | 0.00 | 1.21 | 0.00 | 0.00 | 1.21 |
| FUN_009151 | 5.54 | 5.54 | 5.54 | 0.00 | 0.52 | 0.00 | 0.00 | 0.52 |
| FUN_009222 | 6.57 | 5.02 | 6.57 | 0.00 | 0.52 | 0.35 | 0.35 | 0.52 |
| FUN_009264 | 3.63 | 2.25 | 3.81 | 0.00 | 1.56 | 1.56 | 1.73 | 1.73 |
| FUN_009294 | 2.25 | 2.25 | 2.42 | 0.00 | 0.00 | 1.04 | 1.04 | 0.00 |
| FUN_009350 | 4.67 | 4.67 | 4.84 | 0.00 | 0.52 | 0.00 | 0.00 | 0.52 |
| FUN_009516 | 1.38 | 1.38 | 1.38 | 0.00 | 0.00 | 0.00 | 0.00 | 0.00 |
| FUN_009552 | 5.36 | 6.40 | 5.54 | 0.00 | 1.04 | 1.38 | 1.38 | 1.04 |
| FUN_009576 | 4.67 | 4.33 | 4.84 | 0.00 | 2.77 | 2.60 | 2.60 | 2.94 |
| FUN_009991 | 5.54 | 5.71 | 5.54 | 0.00 | 2.42 | 0.69 | 1.04 | 2.42 |
| FUN_010039 | 3.29 | 3.29 | 3.11 | 0.00 | 1.21 | 1.38 | 1.21 | 1.21 |
| FUN_010054 | 6.92 | 6.92 | 6.57 | 0.00 | 1.38 | 2.08 | 2.25 | 1.38 |
| FUN_010164 | A | A | A | 0.00 | 1.21 | 0.00 | 0.17 | 1.21 |
| FUN_010225 | 9.52 | 9.52 | 9.34 | 0.00 | 0.35 | 0.00 | 0.00 | 0.52 |
| FUN_010331 | 2.08 | 1.73 | 1.90 | 0.00 | 0.52 | 0.52 | 0.52 | 0.52 |
| FUN_010336 | 13.49 | 12.63 | 13.32 | 0.00 | 2.25 | 3.46 | 3.98 | 2.25 |
| FUN_010366 | 4.50 | 4.67 | 4.15 | 0.00 | 0.17 | 0.00 | 0.17 | 0.17 |
| FUN_010414 | 1.90 | 1.73 | 1.90 | 0.00 | 0.35 | 0.52 | 0.52 | 0.35 |
| FUN_010496 | 3.63 | 3.63 | 3.63 | 0.00 | 0.35 | 0.35 | 0.52 | 0.35 |
| FUN_010499 | 1.04 | 1.04 | 0.87 | 0.00 | 0.00 | 0.00 | 0.00 | 0.00 |
| FUN_010529 | 4.15 | 7.09 | 5.36 | 2.42 | 4.15 | 3.46 | 4.15 | 3.11 |
| FUN_010575 | 1.38 | 1.38 | 1.38 | 0.00 | 0.00 | 0.00 | 0.00 | 0.00 |
| FUN_010594 | 3.98 | 3.98 | 3.81 | 0.00 | 0.35 | 0.00 | 0.35 | 0.35 |
| FUN_010651 | 3.46 | 4.15 | 3.46 | 0.17 | 0.69 | 0.00 | 0.00 | 0.87 |
| FUN_010656 | 4.15 | 4.15 | 4.15 | A | 0.69 | 0.00 | 0.00 | 0.69 |
| FUN_010853 | 19.20 | 6.23 | 20.07 | 0.00 | 3.46 | 0.00 | 0.35 | 3.29 |
| FUN_010873 | 7.61 | 6.92 | 7.44 | 0.00 | 2.08 | 0.00 | 0.00 | 2.08 |
| FUN_010899 | 8.82 | 8.48 | 8.13 | 0.00 | 0.52 | 0.17 | 0.00 | 0.52 |
| FUN_010928 | 5.54 | 5.54 | 5.54 | 0.00 | 1.04 | 1.56 | 0.17 | 0.87 |
| FUN_011077 | 5.36 | 5.19 | 5.71 | 0.00 | 0.00 | 0.35 | 0.69 | 0.00 |
| FUN_011081 | 1.56 | 1.90 | 1.90 | 0.00 | 0.52 | 0.17 | 0.17 | 0.52 |
| FUN_011148 | 1.90 | 1.90 | 1.73 | 0.00 | 0.17 | 0.52 | 0.52 | 0.17 |
| FUN_011352 | 3.81 | 3.63 | 4.33 | 0.00 | 0.17 | 0.35 | 0.52 | 0.17 |
| FUN_011359 | 3.46 | 0.17 | 3.29 | 0.17 | 0.17 | 0.35 | 0.35 | 0.35 |
| FUN_011399 | 2.94 | 2.77 | 2.77 | 0.35 | 0.35 | 0.00 | 0.00 | 0.00 |
| FUN_011519 | 4.33 | 3.81 | 2.94 | 0.35 | 0.69 | 0.35 | 0.35 | 0.35 |
| FUN_011522 | 26.12 | 25.95 | 25.95 | 0.00 | 3.81 | 0.87 | 0.35 | 0.52 |
| FUN_011531 | A | A | A | 12.98 | A | 1.90 | 0.17 | 3.81 |
| FUN_011546 | 4.84 | 4.84 | 4.67 | 0.00 | 0.87 | 1.21 | 1.21 | 0.87 |

^*^ While grey boxes labelled with “A” indicate effector gene absence, numbers indicate the SNP ratio (%) in the homolog when compared with the effector gene in R. necatrix strain R18. Green cells indicate identical effector genes that lack SNPs when compared with the sequence in strain R18.

**Supplemental Table 3. Annotated secondary metabolite clusters of *R. necatrix* strain R18.**

| **Contig** | **From** | **To** | **Most similar known cluster** | **Type** | **Organism** | **Similarity** | **Known activity^a^** |
| --- | --- | --- | --- | --- | --- | --- | --- |
| 4 | 28,584 | 63,895 | Swainsonine | Polyketide | *Alternaria oxytropis* | 33% | Phytotoxic (Cook et al., 2017) |
| 5 | 1,431,139 | 1,452,194 | Copalyl diphosphate | Terpene | *Diaporthe amygdali* | 42% | Super-elongation disease in plants (Kawaide, 2006; Rademacher and Graebe, 1979) |
| 5 | 2,494,569 | 2,532,303 | Cytochalasin E/K | NRP+Polyketide | *Aspergillus clavatus* | 23% | Phytotoxic (Sawai et al., 1983; Thomas, 1978) |
| 6 | 409,707 | 450,119 | Wortmanamide A/B | NRP + Polyketide | *Talaromyces wortmannii* | 83% | Unknow (Hai and Tang, 2018) |
| 6 | 1,123,590 | 1,166,755 | Pyriculol | Polyketide | *Neurospora crassa* | 26% | Phytotoxic (Zhao et al., 2019) |
| 6 | 3,495,713 | 3,545,733 | Apicidin | NRP | *Fusarium incarnatum* | 54% | Antiprotozoal activity (Darkin-Rattray et al., 1996) |
| 6 | 3,953,658 | 3,994,676 | Yanuthone D | Polyketide | *Aspergillus niger* | 20% | Antibiotic (Holm et al., 2014) |
| 6 | 4,187,217 | 4,235,557 | Iso-A82775C | Other | *Pestalotiopsis fici* | 41% | Antibacterial activity (Pan et al., 2018) |
| 7 | 2,248,951 | 2,299,047 | Enniatin | NRP | *Fusarium equiseti* | 100% | Phytotoxic (Walton, 1990) |
| 7 | 3,449,415 | 3,495,060 | Naphthalene | Polyketide | *Daldinia eschscholzii* | 33% | Phytotoxic (Xu et al., 2021) |
| 8 | 3,062,127 | 3,105,143 | Melanin | Polyketide | *Glarea lozoyensis* | 100% | No involve in virulence in *R. necatrix* (Shimizu et al., 2014) |
| 9 | 1,168,004 | 1,220,639 | Cytochalasin E/K | NRP+Polyketide | *Aspergillus clavatus* | 30% | Phytotoxic (Sawai et al., 1983; Thomas, 1978) |
| 10 | 2,744,211 | 2,797,043 | Fusaridione A | NRP + Polyketide | *Fusarium heterosporum* | 12% | Unknow (Kalule et al., 2013) |

^a^References:

Cook, D., Donzelli, B. G. G., Creamer, R., Baucom, D. L., Gardner, D. R., Pan, J., Moore, N., Krasnoff, S. B., Jaromczyk, J. W., and Schardl, C. L. (2017). Swainsonine biosynthesis genes in diverse symbiotic and pathogenic fungi. Genes, Genomes, Genetics, 7: 1791-1797.

Kawaide, H. (2006). Biochemical and molecular analyses of gibberellin biosynthesis in fungi. Bioscience, Biotechnology, and Biochemistry, 70: 583-590.

Rademacher, W., and Graebe, J. E. (1979). Gibberellin A4 produced by Sphaceloma manihoticola, the cause of the super elongation disease of cassava (Manihot esculenta). Biochemical and Biophysical Research Communications, 91: 35-40.

Sawai, K., Okuno, T., Fujioka, H., and Furuya, M. (1983). The relation between the phytotoxicity of Cythochalasin E and its molecular structure. Annals of the Phytopathological Society of Japan, 49: 262-265.

Thomas, D. D. (1978). Cytochalasin effects in plants and eukaryotic microbial systems. Frontiers in Biology, 46: 257-275.

Hai, Y., and Tang, Y. (2018). Biosynthesis of long-chain N-acyl amide by a truncated PKS-NRPS hybrid megasynthase in fungi. Journal of the American Chemical Society, 140: 1271-1274.

Zhao, Z., Ying, Y., Hung, Y., and Tang, Y. (2019). Genome mining reveals Neurospora crassa can produce the salicylaldehyde sordarial. Journal of Natural Products, 82: 1029-1033.

Darkin-Rattray, S. J., Gurnett, A. M., Myers, R. W., Dulski, P. M., Crumley, T. M., Allocco, J. J., Cannova, C., Mainke, P. T., Colletti, S. L., Bednarek, M. A., Singh, S. B., Goetz, M. A., Dombrowski, A. W., Polishook, J. D. and Schmatz, D. M. (1996). Apicidin: a novel antiprotozoal agent that inhibits parasite histone deacetylase. Proceedings of the National Academy of Sciences, 93: 13143–13147.

Holm, D. K., Petersen, L. M., Klitgaard, A., Knudsen, P. B., Jarczynska, Z. D., Nielsen, K. F., Gotfredsen, C. H., Larsen, T. O., and Mortensen, U. H. (2014). Molecular and chemical characterization of the biosynthesis of the 6-MSA-derived meroterpenoid yanuthone D in *Aspergillus niger*. Chemistry and Biology, 21: 519-529.

Pan, Y., Liu, L., Guan, F., Li, E., Jin, J., Li, J., Che, Y., and Liu, G. (2018). Characterization of a Prenyltransferase for Iso-A82775C biosynthesis and generation of new congeners of chloropestolides. ACS Chemical Biology, 13: 703-711.

Walton, J. D. (1990). Peptide phytotoxins from plant pathogenic fungi. In Biochemistry of peptide antibiotics. Kleinkauf, H., and von Dohren, H. (eds). Berlin, 179-203 pp.

Xu, D., Xue, M., Shen, Z., Jia, X., Hou, X., Lai, D., and Zhou, L. (2021). Phytotoxic secondary metabolites from fungi. Toxins, 13: 1-65.

Shimizu, T., Tsutae, I., and Kanematsu, S. (2014). Functional analysis of a melanin biosynthetic gene using RNAi-mediated gene silencing in Rosellinia necatrix. Fungal Biology, 118: 413-421.

Thomas, D. D. (1978). Cytochalasin effects in plants and eukaryotic microbial systems. Frontiers in Biology, 46: 257-275.

Kakule, T. B., Sardar, D., Lin, Z., and Schmidt, E. W. (2013). Two related pyrrolidinedione synthetase loci in *Fusarium heterosporum* ATCC 74349 produce divergent metabolites. ACS Chemical Biology, 8: 1549-1557.

**Supplemental Table 4.** **Bacterial strains used in this study.**

| **Strain** | **Species** | **Family** | **Order** | **Medium** | **Gram** |
| --- | --- | --- | --- | --- | --- |
| R102 | *Pseudomonas knackmussii* | Pseudomonadaceae | Pseudomonadales | R2A | Negative |
| R19 | *Pseudomonas corrugata* | Pseudomonadaceae | Pseudomonadales | LBA | Negative |
| R103 | *Bacillus drentensis* | Bacillaceae | Bacillales | R2A | Positive |
| S7 | *Bacillus licheniformis* | Bacillaceae | Bacillales | TSA | Positive |
| R104 | *Paenarthrobacter ureafaciens* | Micrococcaceae | Micrococcales | R2A | Positive |
| S39 | *Arthrobacter enclensis* | Micrococcaceae | Micrococcales | R2A | Positive |
| R109 | *Ochrobactrum intermedium* | Rhizobiaceae | Rhizobiales | R2A | Negative |
| S26 | *Brucella ovis* | Rhizobiaceae | Rhizobiales | TSA | Negative |
| R30 | *Serratia ureilytica* | Enterobacteriaceae | Enterobacterales | TSA | Negative |
| S27 | *Enterobacter soli* | Enterobacteriaceae | Enterobacterales | TSA | Negative |
| R139 | *Microbacterium foliorum* | Microbacteriaceae | Micrococcales | TSA | Positive |
| S23 | *Microbacterium esteraromaticum* | Microbacteriaceae | Micrococcales | TSA | Positive |
| R121 | *Chryseobacterium indoltheticum* | Weeksellaceae | Flavobacteriales | TSA | Negative |
| Ri8 | *Chryseobacterium wanjuense* | Weeksellaceae | Flavobacteriales | TSA | Negative |
| R143 | *Cellulomonas soli* | Cellulomonadaceae | Micrococcales | R2A | Positive |
| R151 | *Paenibacillus lautus* | Paenibacillaceae | Paenibacillales | TSA | Positive |
| S6 | *Paenibacillus illinoisensis* | Paenibacillaceae | Paenibacillales | TSA | Positive |
| R155 | *Achromobacter denitrificans* | Alcaligenaceae | Burkholderiales | LBA | Negative |
| S72 | *Candidimonas bauzanensis* | Alcaligenaceae | Burkholderiales | TSA | Negative |
| R42 | *Solibacillus isronensis* | Planococcaceae | Bacillales | TSA | Positive |
| S15 | *Solibacillus silvestris* | Planococcaceae | Bacillales | LBA | Positive |
| R93 | *Streptomyces flavogriseus* | Streptomycetaceae | Streptomycetales | LBA | Positive |
| Ri17 | *Rhodanobacter spathiphylli* | Rhodanobacteraceae | Xanthomonadales | TSA | Negative |
| Ri21 | *Sphingobium mellinum* | Sphingomonadaceae | Sphingomonadales | TSA | Negative |
| Ri29 | *Pedobacter steynii* | Sphingobacteriaceae | Sphingobacteriales | TSA | Negative |
| Ri32 | *Pedobacter panaciterrae* | Sphingobacteriaceae | Sphingobacteriales | TSA | Negative |
| Ri55 | *Flavobacterium hauense* | Flavobacteriaceae | Flavobacteriales | LBA | Negative |
| Ri56 | *Nocardia coeliaca* | Nocardiaceae | Corynebacteriales | LBA | Positive |
| S13 | *Xanthomonas campestris* | Xanthomonadaceae | Xanthomonadales | R2A | Negative |
| S37 | *Pseudoxanthomonas suwonensis* | Xanthomonadaceae | Xanthomonadales | R2A | Negative |
| S19 | *Brevibacterium sediminis* | Brevibacteriaceae | Micrococcales | LBA | Positive |
| S55 | *Brevibacterium anseongense* | Brevibacteriaceae | Micrococcales | TSA | Positive |
| S25 | *Devosia riboflavina* | Devosiaceae | Rhizobiales | TSA | Negative |
| S29 | *Aeromonas hydrophila* | Aeromonadaceae | Enterobacterales | TSA | Negative |
| S52 | *Kaistia adipata* | Kaistiaceae | Rhizobiales | R2A | Negative |
| S64 | *Exiguobacterium artemiae* | Exiguobacteraceae | Exiguobacterales | TSA | Positive |
| S65 | *Cellulosimicrobium cellulans* | Promicromonosporaceae | Micrococcales | TSA | Positive |
| S71 | *Nocardioides zeae* | Nocardioidaceae | Propionibacteriales | TSA | Positive |
| Si1 | *Herbaspirillum rhizosphaerae* | Oxalobacteraceae | Burkholderiales | TSA | Negative |

**Supplemental Table 5. Primers used in this study.**

| **Primer name** | **Oligonucleotide sequence (5’→3’)** | **Usage** |
| --- | --- | --- |
| RnGAPDH-F | TCAGCGACCAGGAGCTAGTCA | qPCR |
| RnGAPDH-R | TTGAAAAAATTTGGATTGAGCTCTAC | qPCR |
| coRUB-Fw | GAACAGTTTCTCACTGTTGAC | qPCR |
| coRUB-Rv | CGTGAGAACCATAAGTCACC | qPCR |
| RnITS-F | CTGTTCGAGCGTCATTTCAA | qPCR |
| RnITS-R | CCTACCTGATCCGAGGTCAA | qPCR |
| FUN_3304_NdeI_fw | ATCATATGATGATTTCCAACATCCTCCCTGTT | Protein production |
| FUN_3304_BamHI_rv | CCGGATCCTTAGCTGCTGCAGCCACCAAGC | Protein production |
| FUN_4580_NdeI_fw | ATCATATGATGAAGGCAACGATTTTGGACATCG | Protein production |
| FUN_4580_BamHI_rv | CCGGATCCTTATTCGCTGAAGGTGATGGTAACTG | Protein production |
| FUN_5751_NdeI_fw | ATCATATGATGCAGATCTTCACGACAGTCTTGGCCGT | Protein production |
| FUN_5751_BamHI_rv | AGCCGGATCCTCAGCTAGTCCCGGAGCAG | Protein production |
| FUN_9266_NdeI_fw | ATCATATGATGCGTGTCTCAGCTGCTCTCTTCG | Protein production |
| FUN_9266_BamHI_rv | CCGGATCCCTAAAGGGGGTCCGTCCGTC | Protein production |
| FUN_9480_NdeI_fw | ATCATATGATGAAGGCAACTCTGATCTCCGTCGCCGT | Protein production |
| FUN_9480_BamHI_rv | CCGGATCCTTACAGAAGAGCGGCGGCAGCCA | Protein production |

**Supplemental Table 6. Annotation of BLAST and HMMER hits to effector FUN_004580.**

| **Accession ID^a^** | **Annotation^b^** | **Organism^c^** | **Query coverage** | **E value** | **Identity (%)^d^** |
| --- | --- | --- | --- | --- | --- |
| **BLAST** |  |  |  |  |  |
| XM_047979610.1 | Hypothetical protein | *Xylaria bambusicola* | 97% | 0.0 | 71.68% |
| XM_049263194.1 | Hypothetical protein | *Hypoxylon fragiforme* | 96% | 1E-151 | 57.95% |
| XM_051520408.1 | Hypothetical protein | *Durotheca rogersii* | 100% | 1E-144 | 54.59% |
| XM_047931715.1 | Hypothetical protein | *Daldinia caldariorum* | 97% | 2E-141 | 54.34% |
| XM_049244863.1 | Hypothetical protein | *Daldinia decipiens* | 97% | 1E-138 | 53.94% |
| XM_033578433.1 | Hypothetical protein | *Daldinia childiae* | 97% | 4E-137 | 53.06% |
| XM_049303926.1 | Hypothetical protein | *Daldinia loculata* | 97% | 2E-135 | 53.57% |
| XM_047960472.1 | Hypothetical protein | *Annulohypoxylon maeteangense* | 96% | 9E-135 | 51.28% |
| XM_048007749.1 | Hypothetical protein | *Daldinia vernicosa* | 97% | 1E-134 | 53.55% |
| XM_047993846.1 | Hypothetical protein | *Annulohypoxylon truncatum* | 97% | 4E-133 | 52.91% |
| XM_051460929.1 | Hypothetical protein | *Hypoxylon trugodes* | 96% | 9E-132 | 51.79% |
| XM_049311849.1 | Hypothetical protein | *Neoarthrinium moseri* | 100% | 7E-130 | 52.33% |
| XM_007830382.1 | Hypothetical protein | *Pestalotiopsis fici* | 100% | 3E-122 | 49.40% |
| XM_040853841.1 | Hypothetical protein | *Pseudomassariella vexata* | 99% | 4E-120 | 50.61% |
| XM_046099206.1 | Hypothetical protein | *Truncatella angustata* | 83% | 2E-105 | 60.57% |
| XM_046156523.1 | Hypothetical protein | *Microdochium trichocladiopsis* | 95% | 3E-105 | 46.56% |
| XM_018284675.1 | Hypothetical protein | *Pochonia chlamydosporia* | 95% | 2E-102 | 47.31% |
| XM_024896116.1 | Hypothetical protein | *Trichoderma citrinoviride* | 69% | 4E-100 | 56.54% |
| XM_006963277.1 | Hypothetical protein | *Trichoderma reesei* | 69% | 1E-99 | 56.99% |
| XM_024900564.1 | Hypothetical protein | *Trichoderma asperellum* | 96% | 7E-99 | 43.24% |
| XM_038885939.1 | Hypothetical protein | *Colletotrichum karsti* | 67% | 4E-98 | 56.20% |
| XM_018296053.1 | Hypothetical protein | *Colletotrichum higginsianum* | 68% | 1E-97 | 55.40% |
| XM_016786253.1 | Hypothetical protein | *Scedosporium apiospermum* | 94% | 1E-97 | 45.83% |
| XM_036632953.1 | Hypothetical protein | *Colletotrichum siamense* | 61% | 1E-96 | 58.23% |
| XM_045404242.1 | Hypothetical protein | *Colletotrichum gloeosporioides* | 61% | 2E-96 | 58.23% |
| XM_037316359.1 | Hypothetical protein | *Colletotrichum aenigma* | 61% | 2E-96 | 58.23% |
| XM_053175963.1 | Hypothetical protein | *Colletotrichum chrysophilum* | 61% | 2E-96 | 58.23% |
| XM_032030074.1 | Hypothetical protein | *Colletotrichum fructicola* | 61% | 2E-96 | 58.23% |
| XM_049274743.1 | Hypothetical protein | *Colletotrichum spaethianum* | 61% | 4E-96 | 60.00% |
| XM_043142060.1 | Hypothetical protein | *Ustilaginoidea virens* | 68% | 6E-96 | 55.20% |
| XM_046250215.1 | Hypothetical protein | *Ilyonectria robusta* | 67% | 2E-94 | 54.21% |
| XM_018809009.1 | Hypothetical protein | *Trichoderma gamsii* | 69% | 6E-94 | 52.63% |
| XM_044846055.1 | Hypothetical protein | *Fusarium poae* | 95% | 4E-93 | 42.65% |
| XM_007818233.1 | Hypothetical protein | *Metarhizium robertsii* | 95% | 7E-93 | 42.93% |
| XM_024922373.1 | Hypothetical protein | *Trichoderma harzianum* | 69% | 2E-92 | 53.55% |
| XM_036723064.1 | Hypothetical protein | *Colletotrichum truncatum* | 61% | 5E-92 | 57.43% |
| XM_035468007.1 | Hypothetical protein | *Geosmithia morbida* | 67% | 1E-91 | 54.58% |
| XM_014688387.1 | Hypothetical protein | *Metarhizium brunneum* | 95% | 1E-91 | 42.68% |
| XM_009264636.1 | Hypothetical protein | *Fusarium pseudograminearum* | 68% | 2E-91 | 51.97% |
| XM_003651639.1 | Hypothetical protein | *Thermothielavioides terrestris* | 96% | 4E-91 | 43.73% |
| XM_047979610.1 | Hypothetical protein | *Xylaria bambusicola* | 97% | 0.0 | 71.68% |
| XM_049263194.1 | Hypothetical protein | *Hypoxylon fragiforme* | 96% | 1E-151 | 57.95% |
| XM_051520408.1 | Hypothetical protein | *Durotheca rogersii* | 100% | 1E-144 | 54.59% |
| XM_047931715.1 | Hypothetical protein | *Daldinia caldariorum* | 97% | 2E-141 | 54.34% |
| XM_003651639.1 | Hypothetical protein | *Thermothielavioides terrestris* | 96% | 4E-91 | 43.73% |
| XM_025729660.1 | Hypothetical protein | *Fusarium venenatum* | 68% | 1E-90 | 51.25% |
| XM_008095090.1 | Hypothetical protein | *Colletotrichum graminicola* | 61% | 3E-90 | 56.22% |
| XM_014092778.1 | Hypothetical protein | *Trichoderma atroviride* | 69% | 3E-90 | 52.28% |
| XM_049285132.1 | Hypothetical protein | *Colletotrichum lupini* | 60% | 4E-90 | 56.68% |
| CP077948.1 | Hypothetical protein | *Colletotrichum gigasporum* | 92% | 1E-89 | 45.48% |
| XM_053195163.1 | Hypothetical protein | *Colletotrichum fioriniae* | 60% | 6E-89 | 56.56% |
| OW971920.1 | Hypothetical protein | *Trichoderma pseudokoningii* | 67% | 1E-88 | 56.57% |
| XM_031165144.1 | Hypothetical protein | *Fusarium coffeatum* | 68% | 2E-88 | 51.25% |
| XM_006691392.1 | Hypothetical protein | *Thermochaetoides thermophila* | 94% | 7E-88 | 41.45% |
| CP021290.1 | Hypothetical protein | *Trichoderma reesei* | 66% | 1E-87 | 57.88% |
| CP040201.1 | Hypothetical protein | *Trichoderma reesei* | 66% | 1E-87 | 57.88% |
| CP020724.1 | Hypothetical protein | *Trichoderma reesei* | 66% | 1E-87 | 57.88% |
| CP020875.1 | Hypothetical protein | *Trichoderma reesei* | 66% | 1E-87 | 57.88% |
| CP021304.1 | Hypothetical protein | *Trichoderma reesei* | 66% | 1E-87 | 57.88% |
| CP021297.1 | Hypothetical protein | *Trichoderma reesei* | 66% | 1E-87 | 57.88% |
| CP016232.1 | Hypothetical protein | *Trichoderma reesei* | 66% | 1E-87 | 57.88% |
| CP021311.1 | Hypothetical protein | *Trichoderma reesei* | 66% | 1E-87 | 57.88% |
| CP084946.1 | Hypothetical protein | *Trichoderma asperellum* | 94% | 1E-87 | 43.11% |
| XM_022625650.1 | Hypothetical protein | *Colletotrichum orchidophilum* | 68% | 3E-87 | 51.97% |
| XM_011318840.1 | Hypothetical protein | *Fusarium graminearum* | 68% | 4E-87 | 51.97% |
| XM_046123457.1 | Hypothetical protein | *Fusarium flagelliforme* | 68% | 4E-87 | 51.60% |
| CP072834.1 | Hypothetical protein | *Trichoderma asperellum* | 94% | 7E-87 | 42.75% |
| XM_014095116.1 | Hypothetical protein | *Trichoderma virens* | 69% | 1E-86 | 52.13% |
| XM_053146149.1 | Hypothetical protein | *Fusarium falciforme* | 68% | 1E-86 | 50.18% |
| XM_028643441.1 | Hypothetical protein | *Verticillium nonalfalfae* | 61% | 2E-86 | 54.66% |
| XM_003350974.1 | Hypothetical protein | *Sordaria macrospora* | 68% | 3E-86 | 49.64% |
| XM_053050566.1 | Hypothetical protein | *Fusarium keratoplasticum* | 67% | 1E-85 | 50.55% |
| XM_046270538.1 | Hypothetical protein | *Fusarium solani* | 67% | 1E-85 | 50.55% |
| XM_035474047.1 | Hypothetical protein | *Colletotrichum scovillei* | 60% | 1E-85 | 56.68% |
| CP083246.1 | Hypothetical protein | *Epichloe scottii* | 66% | 1E-84 | 53.73% |
| CP101605.1 | Hypothetical protein | *Ustilaginoidea virens* | 65% | 2E-84 | 55.43% |
| CP072756.1 | Hypothetical protein | *Ustilaginoidea virens* | 65% | 2E-84 | 55.43% |
| CP049928.1 | Hypothetical protein | *Ustilaginoidea virens* | 65% | 2E-84 | 55.43% |
| XM_018276255.1 | Hypothetical protein | *Pseudogymnoascus verrucosus* | 93% | 4E-84 | 40.57% |
| CP079833.1 | Hypothetical protein | *Fusarium graminearum* | 68% | 9E-84 | 51.97% |
| XM_003003371.1 | Hypothetical protein | *Verticillium alfalfae* | 61% | 9E-84 | 54.66% |
| CP064806.1 | Hypothetical protein | *Epichloe typhina subsp. clarkii* | 67% | 1E-83 | 51.25% |
| CP003010.1 | Hypothetical protein | *Thermothielavioides terrestris* | 95% | 1E-83 | 43.65% |
| XM_008086879.1 | Hypothetical protein | *Glarea lozoyensis* | 68% | 1E-83 | 49.47% |
| CP117785.1 | Hypothetical protein | *Colletotrichum graminicola* | 65% | 2E-83 | 54.72% |
| CP019475.1 | Hypothetical protein | *Colletotrichum lupini* | 60% | 4E-83 | 56.68% |
| XM_009858316.1 | Hypothetical protein | *Neurospora tetrasperma* | 68% | 4E-83 | 49.28% |
| XM_046198521.1 | Hypothetical protein | *Fusarium redolens* | 69% | 6E-83 | 51.42% |
| CP100344.1 | Hypothetical protein | *Epichloe festucae* | 66% | 1E-82 | 53.36% |
| CP031387.1 | Hypothetical protein | *Epichloe festucae* | 66% | 1E-82 | 53.36% |
| CP098298.1 | Hypothetical protein | *Epichloe typhina subsp. poae* | 67% | 2E-82 | 51.61% |
| CP084938.1 | Hypothetical protein | *Trichoderma atroviride* | 67% | 2E-82 | 52.71% |
| XM_047982779.1 | Hypothetical protein | *Purpureocillium takamizusanense* | 69% | 2E-82 | 48.41% |
| XM_001907097.1 | Hypothetical protein | *Podospora anserina* | 97% | 3E-82 | 37.78% |
| XM_040954205.1 | Hypothetical protein | *Penicillium solitum* | 68% | 3E-82 | 49.82% |
| CP069151.1 | Hypothetical protein | *Verticillium nonalfalfae* | 65% | 4E-82 | 53.61% |
| **HMMER** |  |  |  |  |  |
| A0A5N6KR04_9ROSI | Hypothetical protein | *Carpinus fangiana* | 59,2% | 1.8E-66 | 41,1% |

**^a^**Database accession ID where subject was deposited.

**^b^**Annotation of the best hit using BLAST (tblastn).

**^c^**Organism where the homolog was annotated.

**^d^**Percentage of identity of the query.

**Supplemental Table 7. Annotation of BLAST and HMMER hits to effector FUN_011519.**

| **Accession ID^a^** | **Annotation^b^** | **Organism^c^** | **Query coverage** | **E value** | **Identity (%)^d^** |
| --- | --- | --- | --- | --- | --- |
| **BLAST and HMMER** | | | | | |
| XM_007838424.1 | Hypothetical protein | *Pestalotiopsis fici* | 100% | 6E-10 | 68.57% |
| XM_014319017.1 | Hypothetical protein | *Grosmannia clavigera* | 100% | 1E-09 | 71.43% |
| XM_006668457.1 | Hypothetical protein | *Cordyceps militaris* | 100% | 1E-09 | 71.43% |
| XM_032032777.1 | Hypothetical protein | *Colletotrichum fructicola* | 100% | 2E-09 | 65.71% |
| XM_024655518.1 | Hypothetical protein | *Sordaria macrospora* | 100% | 1E-08 | 68.57% |
| XM_003659946.1 | Hypothetical protein | *Thermothelomyces thermophilus* | 100% | 2E-08 | 71.43% |
| XM_048011148.1 | Hypothetical protein | *Daldinia vernicosa* | 100% | 2E-08 | 62.86% |
| XM_046108154.1 | Hypothetical protein | *Truncatella angustata* | 100% | 6E-08 | 60.00% |
| XM_051511661.1 | Hypothetical protein | *Durotheca rogersii* | 100% | 6E-08 | 62.86% |
| XM_001907105.1 | Hypothetical protein | *Podospora anserina* | 100% | 7E-08 | 65.71% |
| XM_046158086.1 | Hypothetical protein | *Microdochium trichocladiopsis* | 100% | 9E-08 | 60.00% |
| XM_047937750.1 | Hypothetical protein | *Daldinia caldariorum* | 100% | 1E-07 | 60.00% |
| XM_028614903.1 | Hypothetical protein | *Sodiomyces alkalinus* | 100% | 2E-07 | 68.57% |
| XM_051457076.1 | Hypothetical protein | *Hypoxylon trugodes* | 100% | 4E-07 | 65.71% |
| XM_049244688.1 | Hypothetical protein | *Daldinia decipiens* | 100% | 4E-07 | 65.71% |
| XM_014686345.1 | Hypothetical protein | *Metarhizium brunneum* | 100% | 5E-07 | 60.00% |
| XM_003007773.1 | Hypothetical protein | *Verticillium alfalfae* | 100% | 2E-06 | 60.00% |
| XM_003651533.1 | Hypothetical protein | *Thermothielavioides terrestris* | 97% | 3E-11 | 79.41% |
| XM_046265423.1 | Hypothetical protein | *Emericellopsis atlantica* | 97% | 2E-09 | 73.53% |
| XM_008095371.1 | Hypothetical protein | *Colletotrichum graminicola* | 97% | 5E-09 | 70.27% |
| XM_040863558.1 | Hypothetical protein | *Pseudomassariella vexata* | 97% | 7E-09 | 67.65% |
| XM_960199.2 | Hypothetical protein | *Neurospora crassa* | 97% | 2E-08 | 76.47% |
| XM_047972197.1 | Hypothetical protein | *Xylaria bambusicola* | 97% | 4E-08 | 64.71% |
| XM_049302929.1 | Hypothetical protein | *Daldinia loculata* | 97% | 2E-07 | 64.71% |
| XM_049293600.1 | Hypothetical protein | *Colletotrichum lupini* | 97% | 2E-07 | 67.57% |
| XM_035467298.1 | Hypothetical protein | *Geosmithia morbida* | 97% | 4E-07 | 61.76% |
| XM_022620490.1 | Hypothetical protein | *Colletotrichum orchidophilum* | 97% | 4E-07 | 62.16% |
| XM_044863015.1 | Hypothetical protein | *Hirsutella rhossiliensis* | 97% | 7E-07 | 67.65% |
| XM_047991640.1 | Hypothetical protein | *Purpureocillium takamizusanense* | 97% | 1E-06 | 70.59% |
| XM_007808999.1 | Hypothetical protein | *Metarhizium acridum* | 97% | 2E-06 | 61.76% |
| XM_009653120.1 | Hypothetical protein | *Verticillium dahliae* | 97% | 2E-06 | 61.76% |
| XM_046253466.1 | Hypothetical protein | *Ilyonectria robusta* | 97% | 3E-06 | 61.76% |
| XM_053054048.1 | Hypothetical protein | *Fusarium keratoplasticum* | 97% | 5E-05 | 55.88% |
| XM_031161470.1 | Hypothetical protein | *Fusarium coffeatum* | 97% | 6E-05 | 55.88% |
| XM_025733484.1 | Hypothetical protein | *Fusarium venenatum* | 97% | 1E-04 | 55.88% |
| XM_023579291.1 | Hypothetical protein | *Fusarium fujikuroi* | 97% | 2E-04 | 52.94% |
| XM_046198815.1 | Hypothetical protein | *Fusarium redolens* | 97% | 2E-04 | 52.94% |
| XM_053149576.1 | Hypothetical protein | *Fusarium falciforme* | 97% | 3E-04 | 55.88% |
| XM_046283930.1 | Hypothetical protein | *Fusarium solani* | 97% | 4E-04 | 55.88% |
| XM_029896370.1 | Hypothetical protein | *Pyricularia pennisetigena* | 94% | 2E-08 | 72.73% |
| XM_003720841.1 | Hypothetical protein | *Pyricularia oryzae* | 94% | 9E-08 | 69.70% |
| XM_014087113.1 | Hypothetical protein | *Trichoderma atroviride* | 94% | 4E-07 | 63.64% |
| XM_024898021.1 | Hypothetical protein | *Trichoderma citrinoviride* | 94% | 5E-07 | 66.67% |
| XM_031130497.1 | Hypothetical protein | *Pyricularia grisea* | 94% | 1E-06 | 69.70% |
| XM_024548995.1 | Hypothetical protein | *Trichoderma gamsii* | 94% | 3E-06 | 60.61% |
| XM_024901739.1 | Hypothetical protein | *Trichoderma asperellum* | 94% | 8E-06 | 60.61% |
| XM_024919849.1 | Hypothetical protein | *Trichoderma harzianum* | 94% | 1E-05 | 66.67% |
| XM_043141820.1 | Hypothetical protein | *Ustilaginoidea virens* | 91% | 6E-07 | 65.62% |
| XM_009229853.1 | Hypothetical protein | *Gaeumannomyces tritici* | 91% | 5E-06 | 59.38% |
| CP023323.1 | Hypothetical protein | *Cordyceps militaris* | 88% | 2E-07 | 74.19% |
| CP003010.1 | Hypothetical protein | *Thermothielavioides terrestris* | 88% | 3E-07 | 77.42% |
| XM_040822091.1 | Hypothetical protein | *Metarhizium album* | 88% | 2E-06 | 70.97% |
| XM_049264651.1 | Hypothetical protein | *Hypoxylon fragiforme* | 88% | 3E-06 | 58.06% |
| XM_009857960.1 | Hypothetical protein | *Neurospora tetrasperma* | 88% | 4E-06 | 74.19% |
| CP045886.1 | Hypothetical protein | *Beauveria bassiana* | 88% | 4E-06 | 61.29% |
| CP098299.1 | Hypothetical protein | *Epichloe typhina* | 88% | 4E-06 | 67.74% |
| CP098307.1 | Hypothetical protein | *Epichloe typhina* | 88% | 4E-06 | 67.74% |
| CP064797.1 | Hypothetical protein | *Epichloe typhina* | 88% | 5E-06 | 67.74% |
| CP003002.1 | Hypothetical protein | *Thermothelomyces thermophilus* | 88% | 8E-06 | 70.97% |
| CP031390.1 | Hypothetical protein | *Epichloe festucae* | 88% | 2E-05 | 67.74% |
| CP100347.1 | Hypothetical protein | *Epichloe festucae* | 88% | 2E-05 | 67.74% |
| XM_031143259.1 | Hypothetical protein | *Thyridium curvatum* | 88% | 2E-05 | 64.52% |
| CP098267.1 | Hypothetical protein | *Epichloe bromicola* | 88% | 2E-05 | 67.74% |
| CP064805.1 | Hypothetical protein | *Epichloe typhina* | 88% | 2E-05 | 67.74% |
| CP049927.1 | Hypothetical protein | *Ustilaginoidea virens* | 88% | 2E-05 | 64.52% |
| CP072755.1 | Hypothetical protein | *Ustilaginoidea virens* | 88% | 2E-05 | 64.52% |
| CP101604.1 | Hypothetical protein | *Ustilaginoidea virens* | 88% | 2E-05 | 64.52% |
| CP099637.1 | Hypothetical protein | *Epichloe amarillans* | 88% | 2E-05 | 67.74% |
| CP098274.1 | Hypothetical protein | *Epichloe elymi* | 88% | 2E-05 | 67.74% |
| CP077954.1 | Hypothetical protein | *Colletotrichum gigasporum* | 88% | 3E-05 | 64.52% |
| XM_007825388.2 | Hypothetical protein | *Metarhizium robertsii* | 88% | 3E-05 | 61.29% |
| XM_035474232.1 | Hypothetical protein | *Colletotrichum scovillei* | 88% | 4E-05 | 64.71% |
| CP086364.1 | Hypothetical protein | *Purpureocillium takamizusanense* | 88% | 5E-05 | 70.97% |
| XM_040763701.1 | Hypothetical protein | *Sporothrix brasiliensis* | 88% | 6E-05 | 67.74% |
| AB669186.1 | Hypothetical protein | *Colletotrichum orbiculare* | 88% | 7E-05 | 61.29% |
| CP096782.1 | Hypothetical protein | *Nigrospora oryzae* | 88% | 7E-05 | 64.52% |
| CP069148.1 | Hypothetical protein | *Verticillium nonalfalfae* | 88% | 9E-05 | 58.06% |
| CP069139.1 | Hypothetical protein | *Verticillium nonalfalfae* | 88% | 9E-05 | 58.06% |
| CP019480.1 | Hypothetical protein | *Colletotrichum lupini* | 88% | 1E-04 | 64.71% |
| XM_018304892.1 | Hypothetical protein | *Colletotrichum higginsianum* | 88% | 2E-04 | 58.82% |
| CP010982.1 | Hypothetical protein | *Verticillium dahliae* | 88% | 3E-04 | 58.06% |
| CP009079.1 | Hypothetical protein | *Verticillium dahliae* | 88% | 3E-04 | 58.06% |
| CP058936.1 | Hypothetical protein | *Metarhizium brunneum* | 88% | 3E-04 | 61.29% |
| LR026964.1 | Hypothetical protein | *Podospora comata* | 88% | 4E-04 | 64.52% |
| CP071116.1 | Hypothetical protein | *Trichoderma virens* | 85% | 7E-06 | 66.67% |
| XM_014102688.1 | Hypothetical protein | *Trichoderma virens* | 85% | 2E-05 | 66.67% |
| CP084938.1 | Hypothetical protein | *Trichoderma atroviride* | 85% | 4E-05 | 63.33% |
| CP091464.1 | Hypothetical protein | *Pyricularia oryzae* | 85% | 7E-05 | 66.67% |
| CP084944.1 | Hypothetical protein | *Trichoderma asperellum* | 85% | 8E-05 | 60.00% |
| CP072831.1 | Hypothetical protein | *Trichoderma asperellum* | 85% | 1E-04 | 60.00% |
| CP034210.1 | Hypothetical protein | *Pyricularia oryzae* | 85% | 2E-04 | 66.67% |
| CP060336.1 | Hypothetical protein | *Pyricularia oryzae* | 85% | 2E-04 | 66.67% |
| CP099702.1 | Hypothetical protein | *Pyricularia oryzae* | 85% | 2E-04 | 66.67% |
| CP050926.1 | Hypothetical protein | *Pyricularia oryzae* | 85% | 2E-04 | 66.67% |
| CP075865.1 | Hypothetical protein | *Trichoderma simmonsii* | 85% | 2E-04 | 66.67% |
| CP071108.1 | Hypothetical protein | *Trichoderma virens* | 85% | 2E-04 | 66.67% |
| OW971923.1 | Hypothetical protein | *Trichoderma pseudokoningii* | 85% | 3E-04 | 66.67% |
| CP021293.1 | Hypothetical protein | *Trichoderma reesei* | 85% | 3E-04 | 66.67% |
| CP016235.1 | Hypothetical protein | *Trichoderma reesei* | 85% | 3E-04 | 66.67% |
| CP021307.1 | Hypothetical protein | *Trichoderma reesei* | 85% | 3E-04 | 66.67% |

**^a^**Database accession ID where subject was deposited.

**^b^**Annotation of the best hit using BLAST (tblastn).

**^c^**Organism where the homolog was annotated.

**^d^**Percentage of identity of the query.
